# Designing a multi-serotype Dengue virus vaccine: an *in silico* approach to broad-spectrum immunity

**DOI:** 10.1101/2024.12.02.626364

**Authors:** Ankita Singh, Oksana Glushchenko, Alina Ustiugova, Khadija M Alawi, Mikhail Korzinkin, Alex Zhavronkov, Filippo Castiglione

## Abstract

Dengue virus infection represents a major global health issue, with four distinct serotypes complicating the challenge of developing a vaccine due to the need for balanced, long-lasting immunity against all serotypes. Current vaccines have limitations, including an increased risk of severe dengue in seronegative individuals and moderate efficacy, highlighting the need for more effective solutions. Our study aimed to design a multi-serotype Dengue virus vaccine using a computational approach to achieve broad-spectrum immunity. We employed advanced computational tools and algorithms to predict B-cell and T-cell epitopes, ensuring the selection of antigenic targets that provide comprehensive protection against all four serotypes. The methodology included tools for B-cell epitope prediction, tools for MHC class II and I peptide predictions, and tools for toxicity and allergenicity screening to ensure the safety of the vaccine candidates. Our results identified 21 B-cell epitopes, 15 CTL peptides, and 12 HTL peptides, validated for safety regarding toxicity and allergenic potential. The vaccine construct incorporated the adjuvant β-defensin-3 and specific linkers to enhance immunogenicity and stability. Tertiary structure prediction, Ramachandran plot analysis, and stereochemical examination confirmed the stability and quality of the vaccine model.

These findings demonstrate the potential of computational methods in addressing the complex challenges of Dengue virus vaccine development. Our computational approach offers a novel pathway for vaccine design, potentially accelerating the development of effective multi-serotype vaccines. This study provides a promising foundation for future research and clinical validation, marking a significant step forward in dengue vaccine development.

## Introduction

The global burden of Dengue virus (DENV) infections has been rising sharply in recent years, making the development of an effective vaccine a top public health priority. Dengue fever, caused by four distinct serotypes of the dengue virus (DENV-1, DENV-2, DENV-3, and DENV-4), infects millions of people annually and places a heavy strain on healthcare systems, particularly in tropical and subtropical regions (Nasar et al. 2022; Hou et al. 2022). An effective vaccine must provide immunity against all four serotypes to prevent the disease comprehensively (Mustafa & Agrawal 2008; van Eerde et al. 2019). Secondary infection with a different serotype can lead to severe conditions such as dengue hemorrhagic fever and dengue shock syndrome, making it essential for a vaccine to offer balanced protection across all serotypes (Lim et al. 2021).

Despite the urgent need, developing a dengue vaccine faces several significant challenges and limitations. The unique and complex immunopathology of dengue complicates vaccine development (Ghosh & Dar 2015). Additionally, the lack of suitable animal models for the disease and the absence of reliable markers of protective immunity further hinder progress (Wan et al. 2013). The only licensed vaccine, *Dengvaxia*®, has shown limitations, including an increased risk of severe dengue in seronegative individuals and only moderate efficacy. This underscores the need for more effective vaccines capable of protecting against all four serotypes (Anasir & Poh 2022; van Eerde et al. 2019). Collaborative efforts by vaccine manufacturers, regulatory bodies, and policymakers are essential to support vaccine development and standardize field trials, which could make a safe and efficacious dengue vaccine a reality soon (Ghosh & Dar 2015).

Recent advancements in Dengue virus vaccine design have focused on several novel methods, including peptide block entropy analysis, synthetic virology, and deep sequence-coupled biopanning. Peptide block entropy analysis involves calculating the information content of blocks of peptides from a multiple sequence alignment of homologous proteins, providing broad coverage of variant antigens. This method has been applied to the proteomes of the four serotypes of DENV, identifying blocks of peptides that cover 99% of available sequences with five or fewer unique peptides (Olsen et al. 2011). Synthetic virology aims to generate and engineer synthetic viruses, including RNA viruses like DENV, to explore novel approaches for developing vaccines and virus-based therapies (Goz et al. 2018). Deep sequence-coupled biopanning identifies the protein epitopes of antibodies present in human polyclonal serum, providing potential targets for diagnostic and vaccine development (Frietze et al. 2017).

Computational approaches in vaccine design have transformed the field by providing efficient, cost-effective, and rapid alternatives to traditional methods. These techniques use bioinformatics tools to identify potential vaccine candidates, design multi-epitope vaccines, and stimulate immune responses. Computational methods enable the rapid development of potential vaccines in a relatively short time and are recognized as more efficient and economical when compared to traditional laboratory procedures (Huang et al. 2022). These approaches have been successfully used to design multi-epitope vaccines for various diseases, demonstrating promising results. Molecular dynamics (MD) simulations are used to confirm the binding stability and residual flexibility of vaccine constructs, ensuring their stable dynamic and favorable features (Albutti 2022). Despite the advantages, computational approaches have limitations, such as the need for experimental validation to confirm their efficacy and safety in real-world conditions (Chukwudozie et al. 2021).

Broad-spectrum immunity in Dengue virus vaccines is a critical area of research due to the complexity and variability of the virus. One approach involves the development of vaccines that induce pan-serotype neutralizing antibodies and antigen-specific T-cell responses. For instance, a DNA vaccine candidate combining the envelope protein domain III (EDIII) of DENV serotypes 1-4 and a DENV-2 non-structural protein 1 (NS1) protein-coding region has been shown to induce such responses (Sankaradoss et al. 2022). Another strategy involves designing vaccine components as serotype-specific consensus coding sequences derived from different genotypes, ensuring the epitopes are structurally conserved and immunogenic (Sankaradoss et al. 2022). Despite these advancements, challenges such as antibody-dependent enhancement (ADE) complicate the development of broad-spectrum vaccines (He et al. 2021).

In response to these challenges, our study utilized computational approaches to design a multi-serotype Dengue virus vaccine aimed at broad-spectrum immunity. We employed various computational tools and algorithms to predict B-cell and T-cell epitopes, ensuring the selection of antigenic targets that provide comprehensive protection against all four serotypes. Our methodology included the use of BepiPred-2.0 for B-cell epitope prediction, NetMHCII, and NetMHCI for MHC class II and I binding predictions respectively, and tools for toxicity like CSMToxin and allergenicity screening with AllerTop to ensure the safety of the vaccine candidates. Our approach identified 21 B-cell epitopes, 15 CTL peptides, and 12 HTL peptides, further validated for their non-toxicity and non-allergenicity. The vaccine construct included adjuvant β-defensin-3 and specific linkers to enhance immunogenicity and stability. The tertiary structure prediction, Ramachandran plot analysis, and ERRAT stereochemical examination confirmed the quality and stability of the vaccine construct.

## Materials and Methods

For all computational prediction tools, we focused on conserved sequences from the four Dengue serotypes of the three membrane molecules EDIII, prM, and NS1 (Kok et al. 2023; Fahimi et al. 2018).

For each serotype and each membrane molecule EDIII, prM and NS1 the NCBI database records many variants. We downloaded the most prominently reported variants in different regions of the world. The accession numbers (and the country of origin) are given in Table 1-3.

**Table 1.** NCBI identifiers of all DENV Envelope Domain (EDIII) protein variants used in the study.

| Protein | Viral serotype | Accession number | Region |
| --- | --- | --- | --- |
| EDIII | DENV-1 EDIII | ABK54369.1 | China |
|  |  | ABG75766.1 | Hawaii |
|  |  | P33478.2 | Singapore |
|  |  | 6DFJ\|E | United States |
|  |  | 5VIC\|E | Nauru/West Pac |
|  |  | ALG00119.1 | Pakistan |
|  |  | ALG00128.1 | Pakistan |
|  | DENV-2 EDIII | BCG29765.1 | Japan |
|  |  | BBH51307.1 | Bangladesh |
|  |  | P29991.1 | PDK53 |
|  |  | P30026.1 | China |
|  |  | P27914.1 | Tonga |
|  |  | 2JSF_A | Peru |
|  |  | 3J8D | Thailand |
|  |  | AHC72406.1 | Pakistan |
|  |  | AFP56203.1 | Pakistan |
|  |  | AIU39217.1 | Pakistan |
|  |  | P07564.2 | Jamaica |
|  |  | P14338.1 | Malaysia |
|  | DENV-3 EDIII | AMQ36111.1 | France |
|  |  | BAM28865.1 | Japan |
|  |  | QEP41653.1 | Mumbai, India |
|  |  | UBI73841.1 | India |
|  |  | QTE05719.1 | Yangon, Myanmar |
|  |  | Q99D35.1 | China |
|  |  | Q5UB51.1 | Singapore |
|  |  | AHC72431 | Pakistan |
|  |  | AHC72430 | Pakistan |
|  |  | AHC72427 | Pakistan |
|  | DENV-4 EDIII | NP_740317.1 | Maryland |
|  |  | BCG29769.1 | Japan |
|  |  | AEV66314.1 | Vietnam |
|  |  | P09866.2 | Dominica |
|  |  | Q5UCB8.1 | Singapore |
|  |  | AHC72432.1 | Pakistan |

**Table 2.** NCBI identifiers of all DENV prM protein variants used in the study.

| Protein | Viral serotype | Accession number | Region |
| --- | --- | --- | --- |
| prM | DENV-1 prM | ABO21759.1 | East Timor |
|  |  | AAL67810.1 | Mexico |
|  |  | NP_733807.2 | Bethesda, Maryland |
|  |  | NP_722459.2 | Bethesda, Maryland |
|  |  | P27909.2 | Brazil |
|  |  | P27913.1 | Jamaica |
|  | DENV-2 prM | AAL67814.1 | Mexico |
|  |  | ARO34432.1 | Central America |
|  |  | ATQ48916.1 | India |
|  |  | P30026.1 | China |
|  |  | QZZ92461.1 | Saudi Arabia |
|  |  | AGV28546.1 | Pakistan |
|  | DENV-3 prM | AAL67821.1 | Mexico |
|  |  | AFN80339.1 | Australia |
|  |  | AAU29556.1 | Mozambique |
|  |  | AVW83051.1 | Pakistan |
|  | DENV-4 prM | QXF68706.1 | Eastern Uttar Pradesh, India |
|  |  | AAL67829.1 | Mexico |
|  |  | Q5UCB8.1 | Singapore |

**Table 3.** NCBI identifiers of all DENV NS1 protein variants used in the study.

| Protein | Viral Serotype | Ref Accession no | Region |
| --- | --- | --- | --- |
| NS1 | DENV-1 NS1 | P27909.2 | Brazil |
|  |  | P17763.2 | Nauru/West Pac/1974 |
|  |  | CAO01554.1 | Saudi Arabia, Jeddah |
|  |  | BBG62286.1 | northern Vietnam |
|  |  | QHE24281.1 | Philippines |
|  |  | AKK23333.1 | China |
|  | DENV-2 NS1 | P07564.2 | Jamaica |
|  |  | CAA78918.1 | Australia |
|  |  | AAD11533.1 | Japan |
|  |  | AAK67712.1 | Australia |
|  |  | AHC72409.1 | Pakistan |
|  |  | AIU39216.1 | Pakistan |
|  |  | AAA66406.1 | USA |
|  | DENV-3 NS1 | AGW99229.1 | India |
|  |  | AAZ94620.1 | South America |
|  |  | AHC72428.1 | Pakistan |
|  |  | UBI73841.1 | India |
|  | DENV-4 NS1 | AHC72432 | Pakistan |
|  |  | NP_740318 | USA |
|  |  | AUF73704.1 | China |
|  |  | APT69957.1 | Brazil |
|  |  | AUF73704.1 | China |
|  |  | BAJ72475.1 | Philippines |

In the first stage, for each Dengue serotype and each protein ESIII, prM and NS1 we performed a multiple sequence alignment (Madeira et al. 2024; EMBL-EBI n.d.) to identify the conserved sequence ‘chunks’ that are common among the identified variants. This resulted in several conserved regions as reported in Table 4.

**Table 4.**
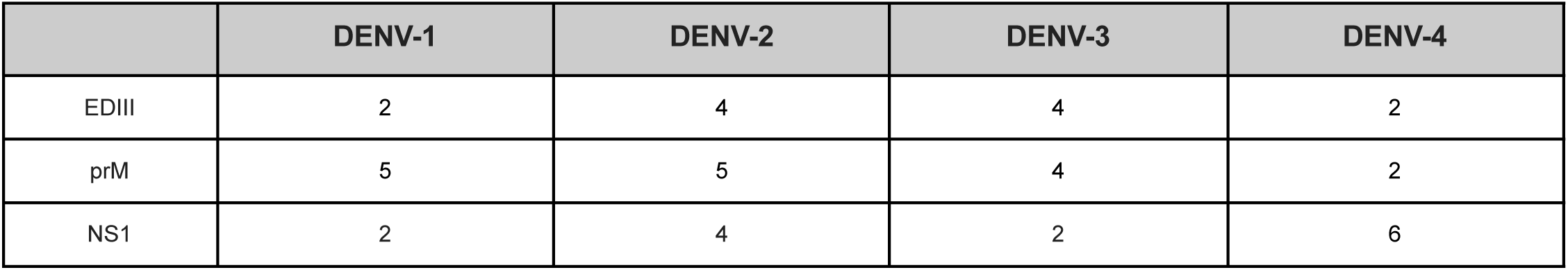
Conserved regions for each protein and each dengue serotype.

For predicting B-cell epitopes, we used the BepiPred-2.0 tool (Jespersen et al. 2017). BepiPred-2.0 is trained on epitope data from crystal structures, enhancing its predictive accuracy. It combines sequence-based features and machine learning techniques to accurately identify potential epitope regions. The epitope threshold was set at 0.6, corresponding to a specificity of 0.95 and a sensitivity of 0.09, to ensure high specificity in the predicted epitopes. The sequences were input into the BepiPred-2.0 server, and the output provided predicted B-cell epitopes based on the chosen threshold.

To maximize the immunogenicity and global applicability of our multi-epitope vaccine, we employed several strategies aimed at enhancing population coverage by selecting peptides that bind to commonly expressed HLA alleles. These strategies ensured that the vaccine candidate could be effective across diverse populations with varying HLA allele distributions (Gupta, Agarwal & Biswas 2018).

To predict cytotoxic T lymphocyte (CTL) epitopes, we utilized NetMHCIpan-4.1 (Reynisson et al. 2020), a cutting-edge tool designed to predict peptide binding to MHC class I molecules. NetMHCIpan-4.1 employs machine learning algorithms trained on experimental binding data and can predict peptide-MHC interactions for any known or unknown MHC class I allele. The peptide sequences derived from Dengue virus proteins were input into the tool, with a specific focus on MHC class I alleles HLA-A02:01 and HLA-B35:01, due to their widespread prevalence in populations endemic to Dengue such as Southeast Asia, Latin America, and the Caribbean (Testa et al. 2012; Malavige et al. 2011). The tool provided binding affinity scores, which were used to classify peptides into strong binders (high affinity) and weak binders (low affinity). Peptides with high affinity were preferred for their potential to induce significant immunological responses.

For predicting HTL peptides, we used the NetMHCIIpan-3.2 tool (Jensen et al. 2018), a well-established method for predicting peptide binding to MHC class II molecules. This tool leverages a pan-specific model that predicts binding affinities for any MHC class II allele, even those without experimental data. The tool provided binding affinity scores indicating how well peptides bind to the selected MHC-II molecules. Peptides were classified based on their binding strength, with high-affinity peptides being selected for their potential to elicit strong immunological responses.

By targeting widely distributed HLA class I and II alleles through these predictive tools, we aimed to enhance the breadth of the vaccine’s efficacy, ensuring it would cover a significant portion of the global population, particularly in regions with high Dengue prevalence. This strategic selection of epitopes was fundamental to maximizing population coverage and vaccine effectiveness across diverse genetic backgrounds.

We used CSM-toxin (Morozov et al. 2023), a computational tool used to predict and analyze cytotoxic sites in protein toxins. CSM-toxin employs machine learning models to identify harmful regions within protein sequences and maps these sites onto 3D structures. The tool was used to screen all predicted epitopes and peptides for toxicity. The input consisted of the peptide sequences, and the output provided insights into the potential cytotoxicity of each peptide, which were crucial for ensuring the safety of the vaccine candidates.

To evaluate the efficacy of our predicted vaccine candidates, we used Vaxigen (Doytchinova & Flower 2007), to predict the antigenicity of selected epitopes and peptides. The tool calculates antigenic scores, with a threshold set at >0.47, and IC_50_ values, with a threshold set at <50. The input consisted of the peptide sequences, and the output provided antigenicity scores, which were used to select the most promising candidates for inclusion in the vaccine construct.

AllerTOP (Dimitrov et al. 2013) is an online server used for *in silico* allergen prediction. It utilizes a machine learning approach based on amino acid E-descriptors, auto-cross covariance transformation, and dataset filtering. The tool was used to calculate the allergenicity scores for each of the predicted B-epitopes, CTL peptides, and HTL peptides. Peptides flagged as allergenic were filtered out to ensure the safety of the vaccine candidates.

The final set of epitopes and peptides were assembled into a multi-epitope vaccine construct. The sequences were joined using specific linkers (i.e., EAAAK, AAY, KK, and GPGPG) and an adjuvant (i.e., β-defensin-3) was added at the N terminus. The tertiary structure of the vaccine construct was predicted using tools such as PSIPRED (Buchan et al. 2013), PDBsum (Laskowski et al. 2018), i-TASSER (Yang & Zhang 2015), Phyre2 (Kelley et al. 2015), RoseTTAFold (Baek et al. 2021), and GalaxyRefine (Heo et al. 2013). The predicted model was evaluated using Ramachandran plot analysis and ERRAT (Colovos & Yeates 1993) stereochemical examination to assess the quality and stability of the structure.

## Results

### Epitope Prediction and Selection

Using our selected Epitope Threshold, we identified 21 predicted peptides against Dengue Virus for all 4 serotypes, see Table 5. These proteins were then analyzed by BepiPred-2.0 and filtered by the Epitope Threshold to yield 21 predicted epitope sequences. BepiPred-2.0 is a computational tool used to predict linear B-cell epitopes, which are essential for understanding immune responses and designing vaccines. For example, BepiPred-2.0 was used to predict linear B-cell epitopes on the envelope (E) and pre-membrane (prM) proteins of the Dengue Virus (DENV), identifying 32 epitopes for the E protein and 17 for the prM protein (Nadugala et al. 2016). This demonstrates the utility of BepiPred-2.0 in predicting B-cell epitopes, crucial for vaccine design and understanding immune responses to Dengue Virus. The peptide sequences derived from Dengue virus proteins were input into the tool, with a specific focus on MHC class I alleles HLA-A02:01 and HLA-B35:01, due to their widespread prevalence in populations endemic to Dengue such as Southeast Asia, Latin America, and the Caribbean (Malavige et al. 2011; Testa et al. 2012).

NetMHCIIpan-3.2 predicts the binding of peptides to MHC class II molecules. Following our approach, we identified 12 unique predicted peptide sequences, see Table 2. This process is vital for designing effective vaccines and understanding immune responses. For instance, one study screened HTL epitopes that could bind strongly to at least 5 human MHC class II alleles for various characteristics, including antigenicity, toxicity, allergenicity, and conservancy, selecting 17 HTL epitopes based on these criteria. These findings highlight the importance of predicting MHC class II binding epitopes for Dengue Virus to enhance vaccine design and understand cross-reactive immune responses. We input peptide sequences derived from Dengue virus proteins into NetMHCIIpan-3.2, specifically focusing on alleles HLA-DRB103:01 and HLA-DRB116:01, which are also prevalent in populations most affected by Dengue virus (Gupta, Agarwal, Kumar, et al. 2018; Guzman et al. 2010).

NetMHC-Ipan is used to predict peptides’ binding affinity to MHC class I molecules. Following our approach, we identified 15 unique predicted peptide sequences, see Table 3. This method is particularly useful given the large number of MHC variants and the costly experimental procedures needed to evaluate individual peptide-MHC interactions (Karosiene et al. 2013). For example, the NetMHCpan method, trained on a large set of quantitative MHC binding data, has demonstrated high performance in predicting binding peptides for non-human primates, indicating its broad allelic coverage beyond human MHC molecules (Hoof et al. 2009). This showcases the utility of predictive algorithms in identifying potential epitopes for vaccine design and immunotherapy.

### Toxicity and Allergenicity Screening

We used the tool Allertop to calculate the allergenicity score for each of the 21 B-epitopes, 15 CTL peptides, and 12 HTL peptides, see Tables 5-7. Peptides flagged as allergenic were filtered out. The AllerTOP v2.0 server is a widely used tool for this purpose, helping predict the allergenicity of vaccine constructs by analyzing protein sequences and identifying potential allergenic proteins. For example, AllerTOP v2.0 was used to predict the allergenicity of a vaccine construct designed for Legionella pneumophila, ensuring that the vaccine candidate does not exert an allergic impact on the host (Mahmoud et al. 2022). This underscores its importance in the early stages of vaccine development to mitigate potential allergic reactions.

**Table 5.**
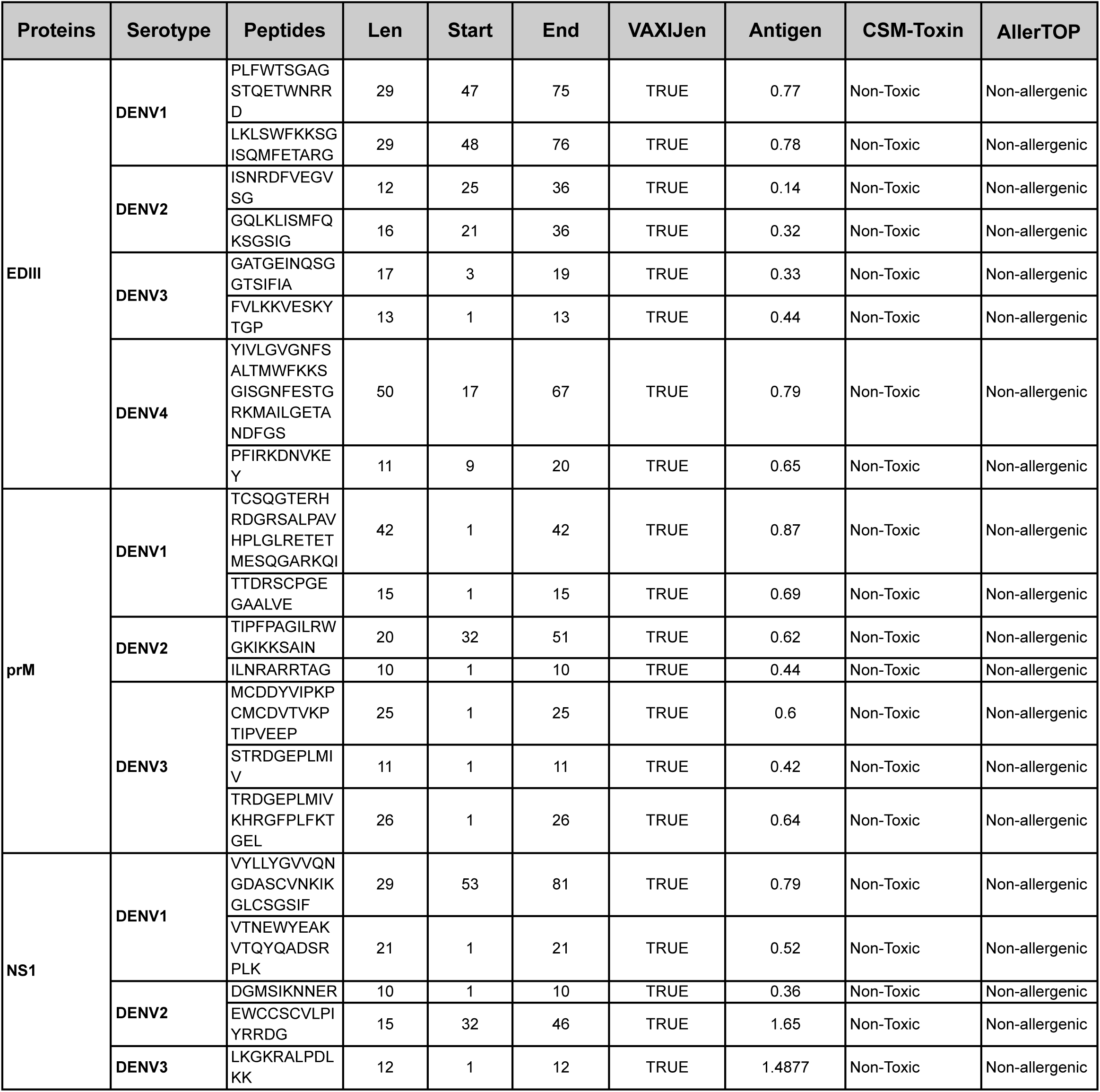
List of predicted B-cell epitopes per protein and serotype.

**Table 6.** Final HTL peptides.

| Protein | Serotype | HTL epitopes | MHC allele | IC50 | Antigenic Score | Allergenicity | Toxicity |
| --- | --- | --- | --- | --- | --- | --- | --- |
| EDIII | DENV1 | ALTGATEINQSGTTTI | HLA- DRB1*07:01 | 34.5 | 0.7846(Antigenic) | Non-allergenic | Non-toxic |
|  | DENV2 | GSYVMCTGSFLFKE | HLA- DRB3*01:01 | 36.8 | 0.8936(Antigenic) | Non-allergenic | Non-toxic |
|  | DENV3 | SNIVIGGDAIKLNIW | HLA- DRB1*13:02 | 21.4 | 0.9058(Antigenic) | Non-allergenic | Non-toxic |
|  | DENV4 | EIRDVNKEVKVGGRI | HLA- DRB1*07:01 | 79.2 | 0.6148(Antigenic) | Non-allergenic | Non-toxic |
| prM | DENV1 | LMLVTPSMAMRCVGI | HLA- DRB1*01:01 | 39.37 | 1.0701(Antigenic) | Non-allergenic | Non-toxic |
|  | DENV2 | KSLLFKTGMNCTLMA | HLA- DRB1*01:01 | 51.4 | 0.47 (Antigenic) | Non-allergenic | Non-toxic |
|  | DENV3 | MCDDYTVYK | HLA- DRB1*03:01 | 1.7 | 0.77(Antigenic) | Non-allergenic | Non-toxic |
|  | DENV4 | DQKAVHADMGWIES | HLA- DRB1*03:01 | 20.5 | 0.6105(Antigenic) | Non-allergenic | Non-toxic |
| NS1 | DENV1 | LTWLGLNSRSTLSM | HLA- DRB3*02:02 | 31.6 | 1.9214(Antigenic) | Non-allergenic | Non-toxic |
|  | DENV2 | AAIKDNRVAHDMGY | HLA- DRB5*01:01, HLA- DRB1*15:01 | 17.24 | 1.1778(Antigenic) | Non-allergenic | Non-toxic |
|  | DENV3 | GSWKLKEASLIEVKT | HLA- DRB1*01:01 | 9.4 | 0.9481(Antigenic) | Non-allergenic | Non-toxic |
|  | DENV4 | DQKAVHADMGWIES | HLA- DRB3*01:01 | 69.4 | 0.6105 (Antigenic) | Non-allergenic | Non toxic |

**Table 7.** Final CTL peptide.

| Protein | Serotype | CTL epitopes | MHC allele | IC50 | Antigenic Score | Allergenicity | Toxicity |
| --- | --- | --- | --- | --- | --- | --- | --- |
| EDIII | DENV1 | KLTIKGVSYV | HLA-A*02:03 | 8.67 | 1.68(Antigenic) | Non-allergenic | Non-toxin |
|  | DENV2 | AMHTALATGA | HLA-A*23:01 | 10.66 | 0.42(Antigenic) | Non-allergenic | Non-toxin |
|  |  | KLTIKGVSYV | HLA-A*68:01 | 38.18 | 1.20(Antigenic) | Non-allergenic | Non-toxin |
|  |  | LSVSLVLGV | HLA-A*68:02 | 42.7 | 1.38(Antigenic) | Non-allergenic | Non-toxin |
|  | DENV3 | TILIKVEPK | HLA-A*11:01 | 46.81 | 1.79(Antigenic) | Non-allergenic | Non-toxin |
|  | DENV4 | RIKGMSYTM | HLA-A*30:01 | 36.8 | 0.73(Antigenic) | Non-allergenic | Non-toxin |
|  |  | KLRIKGVMSY | HLA-A*30:01 | 22.6 | 1.48(Antigenic) | Non-allergenic | Non-toxin |
| prM | DENV1 | SMAMRCVGI | HLA-A*02:06 | 39.69 | 1.5822(Antigenic) | Non-allergenic | Non-toxin |
|  | DENV2 | RQEKGKSLLF | HLA-B*15:01 | 61.97 | 0.4506(Antigenic) | Non-allergenic | Non-toxin |
|  | DENV3 | LTSRGDEPR | HLA-A*68:01 | 37.64 | 1.5125(Antigenic) | Non-allergenic | Non-toxin |
|  | DENV4 | LLFKTTEGI | HLA-A*02:03 | 17.81 | 0.6668(Antigenic) | Non-allergenic | Non-toxin |
| NS1 | DENV1 | LSMTCIAVGV | HLA-B*58:01,<br>HLA-A*02:01 | 21.58 | 2.366(Antigenic) | Non-allergenic | Non-toxin |
|  | DENV2 | AAEGINYAD | HLA-A*02:03,<br>HLA-B*35:01 | 10.45 | 1.1155(Antigenic) | Non-allergenic | Non-toxin |
|  | DENV3 | TAGPHWLQK | HLA-A*11:01 | 49.93 | 1.1355(Antigenic) | Non-allergenic | Non-toxin |
|  | DENV4 | TQTPGWHLGK | HLA-A*31:01 | 48.38 | 1.1272(Antigenic) | Non-allergenic | Non-toxin |

We screened all epitopes and peptides for toxicity using CSM-toxin and filtered out those labeled as toxic. The final set of epitopes and peptides are reported in Tables 5-7. The CSM-Toxin method is a valuable tool for toxicity assessment in vaccine design, accurately identifying peptides and proteins with potential toxicity. This performance highlights the robust and generalizable nature of the model, making it a significant asset in minimizing potential toxicity in the biological development pipeline (Morozov et al. 2023).

### Multi-Epitope Vaccine Construction

The adjuvant β-defensin is included in the construct at the N terminus. Specific linkers (EAAAK, AAY, KK, and GPGPG) are incorporated to differentiate particular epitopes. Adjuvants and linkers play a crucial role in the design of multi-epitope vaccines by enhancing immunogenicity and ensuring the proper presentation of epitopes. For example, a cholera toxin-derived adjuvant was integrated with prioritized T-cell epitopes to design a multi-epitope vaccine, confirmed through molecular docking analysis showing strong structural associations with HLA alleles (Dar et al. 2022). This approach has been successfully applied in various studies to develop vaccines against different pathogens.

Using appropriate linkers with established immunomodulatory activity and stability (EAAAK, KK, AAY, and GPGPG), the various components were combined to create the construct (cfr. Figure 1). Since the β-defensin-3 adjuvant is separated from the epitopes by the linker EAAAK, the construct remains stiff, preserving both the unique functions and structural stability of each epitope. To guarantee humoral and cellular responses, the B-cell epitope was linked with the flexible linker KK, while the CTL and HTL epitopes were linked with the linkers AAY and GPGPG, respectively. This allowed for optimal presentation of the epitopes to the immune system. An improved immune response to correct folding, processing, and presentation of every epitope type ought to be the outcome of such a modular architecture.

**Figure 1.**
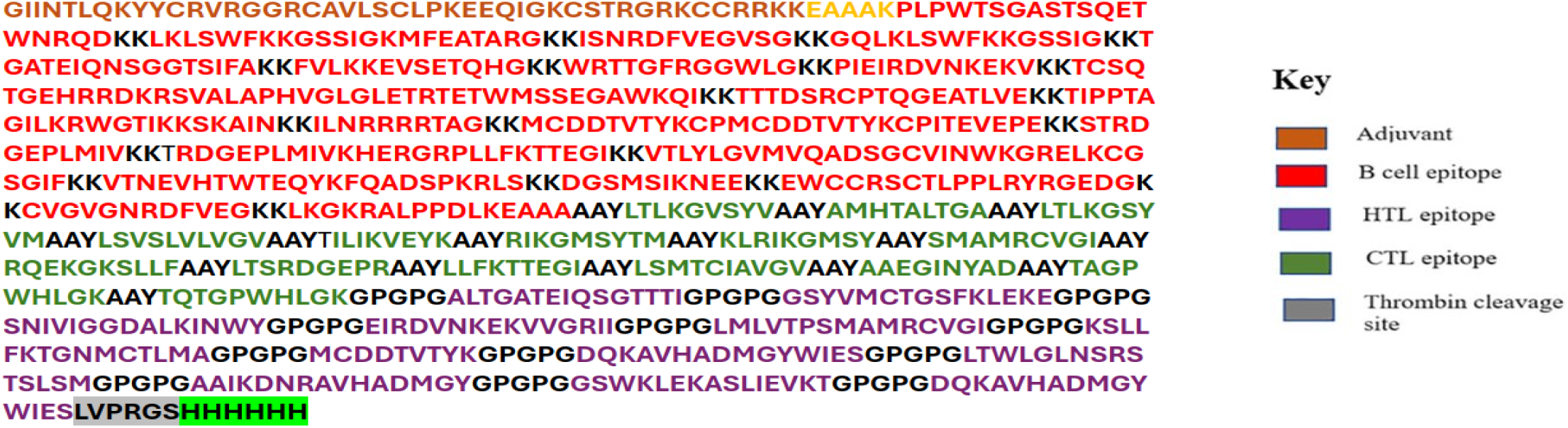
The following list contains every amino acid that is present in the vaccine design. The adjuvant β-defensin is included into the construct at the N terminus. To differentiate particular epitopes, specific linkers (EAAAK, AAY, KK, and GPGPG) are incorporated.

### Vaccine Stability and Optimization

We provide a prediction model for vaccine construction, see Figure 2A. The vaccine’s tertiary structure includes the adjuvant (brown), thrombin site (red), EAAAK linker (yellow), KK linker (dark magenta), AAY linker (lime green), and GGGS linker (cyan); light brown represents all epitopes in the vaccine build. Tertiary structure prediction is crucial for understanding the three-dimensional conformation of the vaccine candidate, essential for its interaction with immune receptors and subsequent immune response. For example, in the design of a candidate vaccine against dengue virus serotypes, researchers employed immunoinformatics and bioinformatics tools to predict and validate the tertiary structure of the vaccine construct (Shoushtari et al. 2024). This ensures high-quality modeling for all vaccine constructs.

**Figure 2.**
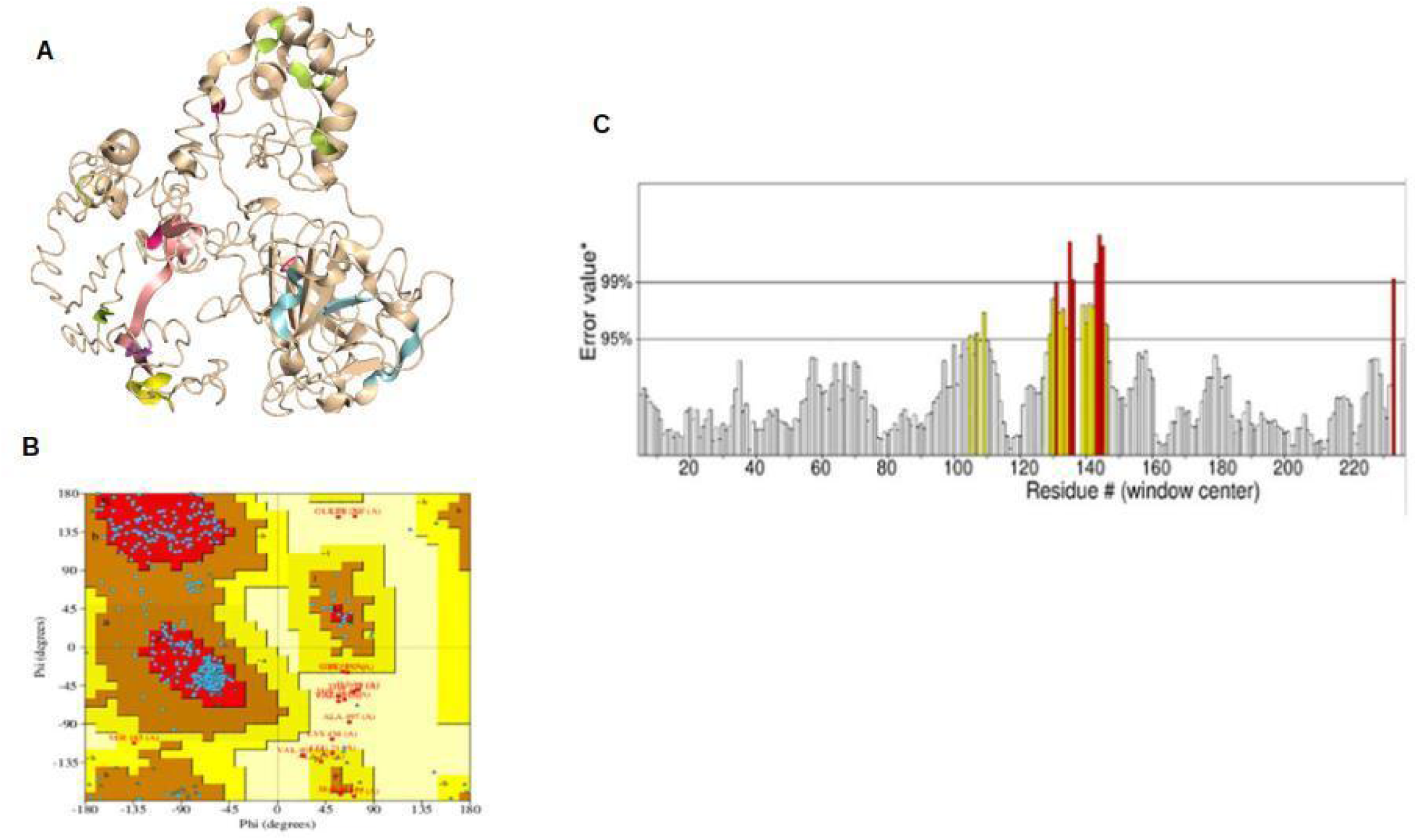
Prediction model for vaccine construction. (A) The vaccine’s tertiary structure. adjuvant (brown), thrombin site (red), EAAAK linker (yellow), KK linker (dark magenta), AAY linker (lime green), and GGGS linker (cyan); light brown represents all epitopes in the vaccine build. (B) The Ramachandran plot analysis for the model’s quality assessment revealed that 84.4% of the residues are present in the most favored regions, with 12% present in additional authorized regions. (C) ERRAT’s stereochemical examination revealed a higher ERRAT value of 93.84, indicating a highly refined structure. In a protein model, the X-axis shows the number of amino acids (#), while the Y-axis reflects the error value*.

The Ramachandran plot analysis for the model’s quality assessment revealed that 84.4% of the residues are present in the most favored regions, with 12% present in additional authorized regions, see Figure 2B. The Ramachandran plot is a crucial tool for assessing the quality of vaccine models by evaluating the conformational angles of amino acid residues in protein structures. For example, the refinement process of a 3D model structure was performed to improve its quality, and the validation was carried out using the Ramachandran plot and ProSA z-score (Touhidinia et al. 2021). This ensures the structural reliability and potential efficacy of vaccine candidates.

ERRAT’s stereochemical examination revealed a higher ERRAT value of 93.84, indicating a highly refined structure, see Figure 2C. In a protein model, the X-axis shows the number of amino acids, while the Y-axis reflects the error value. ERRAT is a widely used tool for assessing the quality of protein structures by analyzing the statistics of non-bonded interactions between different atom types. For example, the ERRAT overall quality factor for the NS1 protein was calculated to be 86.79, indicating a high-quality model (Mukhtar et al. 2022). This highlights the importance of ERRAT in evaluating the structural integrity of vaccine models.

## Discussion

The *in silico* approach for vaccine design marks a notable advancement in vaccinology, utilizing computational tools to streamline and improve the vaccine development process. This method stands out due to several key elements. Firstly, in-silico techniques aid in identifying T-cell and B-cell epitopes, which are essential for vaccine effectiveness. For example, computational prediction tools have been used to pinpoint major histocompatibility complex (MHC) class I and II T-cell epitopes and B-cell epitopes for the Lassa virus glycoprotein. Secondly, these techniques speed up the vaccine development process and reduce costs, making it quicker and more economical compared to traditional methods (García-Machorro et al. 2022). Thirdly, advancements in computational vaccinology have enabled rational vaccine design, grounded in a deep understanding of the pathogen’s biology and the host’s immune response. This approach results in vaccines with higher efficacy and safety (Söllner et al. 2010; Leclerc 2007). Moreover, synthetic biology has provided algorithms for the rapid in-silico identification of relevant protein candidates and the design of new immunogens with enhanced expression, safety, and immunogenicity profiles (Kindsmüller & Wagner 2011). Additionally, in-silico studies support vaccine development in both oncology and infectious diseases, offering a thorough analysis of potential vaccine candidates (Saldanha et al. 2023). Lastly, while in-silico methods are powerful, they are often supplemented by experimental validation to confirm the immunogenicity and protective capabilities of the identified vaccine candidates (Robleda-Castillo et al. 2021). Overall, the innovation of the in-silico approach lies in its ability to integrate vast amounts of biological data, predict immune responses, and streamline the vaccine development process, ultimately leading to more effective and safer vaccines.

The global burden of Dengue virus infections has been rising sharply in recent years. The virus is prevalent in subtropical and tropical areas, infecting up to 400 million people each year, with around 500,000 individuals developing life-threatening conditions (Scaturro et al. 2014). In India, which is considered a dengue hyper-endemic country, the burden of dengue virus infections has also increased. A study conducted in Hyderabad, India, in 2014, highlighted the simultaneous circulation of all four dengue serotypes and co-infections, with a high percentage of severe dengue cases (Vaddadi et al. 2017). In Africa, the burden of dengue is not well documented, but recent studies indicate a high prevalence and significant risk factors associated with the disease. A review of literature published between 1960 and 2020 found that the prevalence of dengue was 29% during outbreak periods and 3% during non-outbreak periods. The leading risk factors included old age, lack of mosquito control, urban residence, climate change, and recent travel history (Mwanyika et al. 2021). The dominant serotypes in Africa were DENV-1 and DENV-2, contributing to 60% of the epidemics (Mwanyika et al. 2021). In Gabon, Central Africa, a surveillance study revealed the recent re-emergence of DENV and highlighted the different potential risks and virus-specific circulation patterns (Ushijima et al. 2021). The overall seroprevalence of DENV during 2014-2017 was estimated to be 20.4% (Ushijima et al. 2021). The increasing global burden of dengue highlights the need for innovative methods to understand and control arbovirus infections. Metabolomics, for example, has been applied to detect alterations in host physiology during infection, providing insights into host-virus dynamics and potential therapeutic targets (Byers et al. 2019). In summary, the global burden of dengue virus infections is substantial and increasing, with significant impacts on public health and economies worldwide. Effective surveillance, innovative diagnostic methods, and targeted interventions are crucial to mitigate the impact of this disease.

The need for a multi-serotype Dengue virus vaccine is driven by several critical factors. Dengue fever is caused by four distinct serotypes of the virus, and infection with one serotype does not confer immunity against the others. Subsequent infections with different serotypes can lead to more severe forms of the disease due to a phenomenon known as antibody-dependent enhancement (Sariol & White 2014). A major challenge in developing a dengue vaccine is the need to induce balanced, long-lasting tetravalent immune responses against all four serotypes. This is crucial because primary infection by any one serotype may predispose individuals to more severe diseases during a heterotypic secondary infection (Sun et al. 2017). The only licensed dengue vaccine, Dengvaxia®, has shown suboptimal protection against some serotypes and has paradoxically increased the risk of severe dengue in seronegative individuals (Hou et al. 2020; Anasir & Poh 2022). The development of a tetravalent vaccine is further complicated by the need to ensure that the vaccine components do not interfere with each other, leading to unbalanced immune responses. Clinical trials have shown that unbalanced replication and immunodominance of one vaccine component over others can result in low efficacy and vaccine-enhanced severe disease (Nivarthi et al. 2021). Given the global distribution of dengue and the co-circulation of all four serotypes in endemic areas, a vaccine that provides balanced protection against all serotypes is essential to reduce morbidity and mortality effectively. Current efforts are focused on developing more effective and affordable vaccines that can provide long-lasting immunity against all four serotypes (van Eerde et al. 2019; Mustafa & Agrawal 2008). In summary, the need for a multi-serotype dengue virus vaccine is underscored by the necessity to provide comprehensive protection against all four serotypes to prevent severe disease and manage the global burden of dengue effectively (Raviprakash et al. 2008; Ali et al. 2022).

Broad-spectrum immunity for Dengue virus can be achieved through various mechanisms, including the use of genetically modified antibodies, innate immune stimulatory agents, and the siRNA pathway. Research has demonstrated that the pathogenesis of severe DENV infection can be inhibited by the therapeutic administration of genetically modified antibodies or RIG-I receptor agonists that stimulate innate immunity. These treatments have shown effectiveness in a more immunocompetent mouse model of DENV infection, which recapitulates many characteristics of severe human disease, such as vascular leakage, hemoconcentration, thrombocytopenia, and liver injury (Pinto et al. 2015). The siRNA pathway plays a central role in the control of viral infections in mosquitoes. Overexpression of Dicer2 (Dcr2) or R2d2 in mosquitoes has been shown to result in the accumulation of 21-nucleotide viral sequences, significantly suppressing the replication of DENV, Zika virus (ZIKV), and chikungunya virus (CHIKV). This indicates a broad-spectrum antiviral response mediated by the siRNA pathway, which can be applied to the development of novel arbovirus control strategies (Dong et al. 2022). The RNAi pathway, particularly the Dicer-2 (dcr2) gene, is a major antiviral component of innate immunity in invertebrates. Polymorphisms in the dcr2 gene have been associated with isolate-specific resistance to DENV in mosquitoes, suggesting that host-pathogen compatibility depends on genotype-by-genotype interactions between dcr2 and the viral genome (Lambrechts et al. 2013). These findings highlight the potential of broad-spectrum antiviral strategies that leverage both the adaptive and innate immune responses to combat DENV and other related viral infections.

Computational tools and algorithms play a crucial role in modern vaccine design by enabling rapid identification and analysis of potential vaccine candidates. These tools utilize bioinformatics and machine learning techniques to predict epitopes, which are the parts of antigens that are recognized by the immune system. Tools like EpiMatrix, Conservatrix, BlastiMer, and Patent-Blast are used to identify T cell epitopes and analyze protein sequences for highly conserved regions and homology with other known proteins. These tools enable researchers to move rapidly from genome sequence to vaccine design by narrowing down the list of putative epitopes to be tested in vitro (De Groot et al. 2001). The performance of T cell epitope prediction tools has been evaluated using datasets that systematically define T cell epitopes recognized in vaccinia virus (VACV) infected mice. These evaluations found that neural network-based predictions trained on both MHC binding and MHC ligand elution data (e.g., NetMHCPan-4.0 and MHCFlurry) achieved the best performance (Paul et al. 2020). Tools like Predivac-2.0 are particularly suited for epitope-based vaccine design against virus-related emerging infectious diseases. This tool maps putative CD4+ T-cell epitopes on the surface glycoproteins of pathogens and is validated for identifying promiscuous and immunodominant CD4+ T-cell epitopes from HIV protein Gag (Oyarzun et al. 2015). Computational experiments are essential for vaccine development as they allow for the analysis of virus particles and immune system reactions before in vitro testing. For example, *in silico* analysis of Papaya mosaic Virus-Like Particles (VLPs) helped clarify particle structure and immune response potential, demonstrating that these nanoparticles can trigger an immune response without fusion with any foreign antigen (Zamani-Babgohari et al. 2019). The benchmarking of epitope prediction tools provides guidance for immunologists on which methods to use and what success rates to expect. This benchmarking is implemented in an open and reproducible format, offering a framework for future comparisons against new tools (Paul et al. 2020). In summary, computational tools and algorithms significantly enhance the efficiency and accuracy of vaccine design by predicting epitopes, evaluating performance, and providing frameworks for future research. These advancements are crucial for developing vaccines against both existing and emerging infectious diseases.

Validation of in-silico vaccine predictions is a crucial step to ensure that the theoretical and computational findings translate into effective and safe vaccines in real-world scenarios. The predictions made by in-silico tools need to be validated through various immunological assays. This includes testing the vaccine constructs in animal models to confirm their efficacy and safety (Karkashan 2024). To understand the immune response elicited by the vaccine construct among different races, population-wise studies can be conducted. This helps in assessing the vaccine’s effectiveness across diverse genetic backgrounds. Molecular dynamics (MD) simulations are used to validate the stability and interaction of the vaccine construct with target receptors, such as Toll-like receptors (TLRs). These simulations help in predicting the stability of the vaccine-receptor complex over time (Nayak et al. 2024; Ghaffar et al. 2024). Computational immune simulations can predict the potential immune response elicited by the vaccine. These simulations provide insights into the types of immunoglobulins and cellular responses that the vaccine might induce (Karkashan 2024). The secondary and tertiary structures of the vaccine constructs are assessed using tools like PSIPRED and SOPMA. These tools predict the structural stability and composition quality of the vaccine, which are essential for its effectiveness (Nayak et al. 2024). The vaccine constructs are evaluated for their allergenicity and antigenicity using servers like AllerTOP, Allergen FP, VaxiJen, and ANTIGENPro. This ensures that the vaccine is safe and capable of eliciting a strong immune response (Nayak et al. 2024). Codon optimization and in-silico cloning are performed to enhance the expression of the vaccine in host cells. This step is followed by computational immune assays to predict the immune response (Nayak et al. 2024; Karkashan 2024).

## Conclusions

Our study introduces a new computational method for designing a multi-serotype Dengue virus vaccine aimed at achieving broad-spectrum immunity. By using computational tools and algorithms, we identified potential B-cell and T-cell epitopes that can protect against all four Dengue virus serotypes. This method not only speeds up the vaccine development process but also offers a cost-effective and efficient alternative to traditional methods, addressing a pressing need in the field of dengue vaccine research.

The main advantage of our study is its ability to quickly identify and validate vaccine candidates using computational methods. This greatly reduces the time and resources needed for vaccine development. Furthermore, our method ensures the selection of antigenic targets that are likely to induce strong and balanced immune responses, which is essential for preventing severe dengue caused by secondary infections with different serotypes. However, the study also has limitations. The computational predictions need to be validated through extensive experimental and clinical trials to confirm their efficacy and safety in real-world scenarios. Additionally, the complexity of the dengue virus and the phenomenon of antibody-dependent enhancement (ADE) pose challenges that require careful consideration in vaccine design.

To improve the study, future research should focus on integrating more advanced computational models that can simulate the immune response in greater detail. Also, incorporating real-world data from diverse populations can help refine the predictions and ensure the vaccine’s effectiveness across different genetic backgrounds. Experimental validation, including animal studies and clinical trials, is essential to confirm the computational findings and address any potential safety concerns. By combining computational and experimental approaches, we can develop a more effective and reliable multi-serotype Dengue virus vaccine, ultimately contributing to better disease control and prevention.

## Author Contributions

AS conducted and coordinated *in silico* studies and contributed to the writing of the manuscript. OG coordinated *in silico* studies and writing, KMA, AU and MK contributed to the writing of the manuscript. AZ and FC supervised the project and provided suggestions to the design and improvement of the manuscript. FC contributed to writing the manuscript and is the corresponding author.

A preliminary draft for the current manuscript was generated with the assistance of DORA (Draft Outline Research Assistant (https://pharma.ai/dora), Insilico Medicine’s LLM-based text drafting assistant. DORA is designed to streamline the process of text drafting, making it faster and simpler. The text generation is curated by over 30 AI agents, powered by Large Language Models, and integrated with internal and other curated databases, to assist in generating high-quality scientific document drafts. Each agent employs Retrieval-Augmented Generation (RAG) to perform comprehensive data collection and analysis, reduce the probability of hallucinations, and provide relevant PubMed links to make the generation of the article more transparent. The draft was further manually curated, substantially extended, and reviewed by all authors.

## Data availability

All data supporting the conclusions of the paper are available in the article and corresponding figures. Tools used in the paper are described in the Materials and Methods section.

